# The cytoplasmic lattice in mammalian eggs sequesters ubiquitination machinery and tubulin in reserve

**DOI:** 10.64898/2026.03.30.715190

**Authors:** Yujie Li, Wei Zheng, Jiyeon Leem, Chunxiang Wu, Shaogeng Tang, Binyam Mogessie, Yong Xiong

## Abstract

The cytoplasmic lattice (CPL) in mammalian eggs is essential for early embryonic development, but its molecular components, structural organization, and functional capacity have remained elusive. Here, using cryo-electron microscopy, we show that the CPL filament in mouse eggs contains repeating units with a periodicity of ∼37 nm, and determine its high-resolution, native structure and complete subunit composition. The CPL architecture organizes maternal-effect proteins, ubiquitination machinery, and tubulin into a highly structured reservoir. Maternal-effect proteins form the scaffold of the CPL to sequester a UBE2D–UHRF1 E2-E3 ubiquitination module and three distinct FBXW E3 ubiquitin ligases, notably all in activity-excluded states. The CPL further contains an αβ-tubulin heterodimer in a GTP-bound state with a calcium ion coordinated to α-tubulin, suggesting microtubule assembly-competent tubulin held in reserve. The CPL structure is capped at each end by a terminal unit that lacks a PADI6 dimer, a scaffold component, thereby preventing further oligomerization. Interactions between maternal-effect proteins in adjacent CPL units promote the assembly of a three-dimensional lattice in the egg cytoplasm. Taken together, our work defines how CPL assembly and architecture prime mammalian eggs for ubiquitin-mediated protein degradation and cytoskeletal remodeling during the egg-to-embryo transition.

## Introduction

How cells efficiently organize their interior to support function is a central problem in biology. This challenge is especially critical in large, long-lived cells, where diffusion limits, spatial constraints, and extended cellular lifespan impose demands beyond those of typical somatic cells^1–3^. Mammalian oocytes are an extreme example. They are among the largest cells in the body and can remain arrested for long periods while maintaining the ability to rapidly undergo meiotic divisions and to support early embryonic development^4,5^. How such cells maintain cytoplasmic order, preserve essential molecular activities in a poised state, and prevent premature activation of developmental programs remains poorly understood.

Classical models of cellular organization, primarily based on small, rapidly dividing cells, focus on membrane-bound organelles and dynamic cytoskeletal networks. These models do not readily explain how oocytes manage the storage, protection, and timely use of vital components over extended timescales. This gap in understanding limits mechanistic insights into key processes that affect reproductive health, such as the causes of age-related decline in oocyte quality and the high rate of aneuploidy in human eggs.

A prominent, but poorly understood feature of mammalian oocytes is the cytoplasmic lattice (CPL), a dense fibrillar network that permeates the cytoplasm and is crucial for the egg-to-embryo transition^6–11^. Loss of CPL components, including the maternal-effect proteins PADI6, NLRP5, TLE6, and OOEP^8,9,12,13^, results in early embryonic arrest, suggesting its functional importance. However, the molecular architecture of the CPL and the principles by which it contributes to cellular organization and functions are only starting to emerge^14,15^.

The CPL has long been proposed to function as a storage site for maternal proteins supporting early embryonic development^14^, yet its structure at the final stage of oocyte maturation remains unresolved. Although the organization of CPLs in germinal vesicle (GV)-stage oocytes has been partially defined^15^ and CPLs persist in metaphase II (MII) eggs^14^, their architecture immediately preceding fertilization has not been established. Resolving this state is important for understanding how the CPL may package and organize essential factors in preparation for embryogenesis. However, the structural complexity of the CPL also presents a major technical obstacle, as conventional biochemical isolation approaches are likely to perturb its native architecture, disrupt inter-filamental contacts, and dissociate more weakly associated factors.

To define the final structural state of the CPL before embryogenesis, we determined its native structure in fertilizable MII eggs. To preserve native CPL architecture, zona pellucida (ZP)-free eggs were deposited onto cryo-electron microscopy (cryo-EM) grids and rapidly ruptured mechanically to generate a thin cytoplasmic layer suitable for imaging. Using single-particle cryo-EM, we determined the structure of the complete CPL building block at 3.5 Å resolution, defined its assembly principles, and gained insight into its potential function. The CPL is formed by translational polymerization of repeating units with a core composed of maternal-effect proteins, ubiquitination machinery, and αβ-tubulin heterodimers. Notably, we found that a UBE2D–UHRF1 ubiquitination module and three distinct FBXW E3 ubiquitin ligases are sequestered in their inhibitory states, along with microtubule assembly-competent, GTP-bound tubulin heterodimers containing a coordinated calcium ion. Preservation of intra-lattice interactions in our preparations allowed us to define how maternal scaffold proteins drive CPL fiber assembly. Together, these findings establish the CPL as a structured maternal reservoir of proteostatic and cytoskeletal resources for early embryonic development.

## Results

### Native structure of the CPL in metaphase II mouse eggs

To understand the ultrastructural organization of the CPL in fully mature eggs, we first characterized mouse eggs by negative-stain transmission electron microscopy (TEM). In our TEM micrographs of ultrathin sections, we consistently observed highly abundant CPL structures broadly distributed throughout the cytoplasm. Morphologically, these assemblies constitute a highly ordered fibrous network composed of stacked, lattice-like filament arrays (Extended Data Fig. 1a). Quantification of individual CPL units within these arrays demonstrates a wide distribution of lengths, with a median of approximately 340 nm and some long filaments extending beyond 1.2 μm (Extended Data Fig. 1b). This macroscopic architecture and the filament length distribution in MII eggs are highly analogous to the lattice structures previously observed in immature oocytes^14^. These observations confirm that the extensive CPL framework is robustly maintained during meiotic maturation, persisting as a dominant ultrastructural feature in the mature egg.

To further resolve the molecular structure of CPL in mature eggs at high resolution, we developed a purification-free cryo-EM pipeline to visualize the endogenous CPL in the native context of MII eggs. As shown in Fig.1a, ZP-free MII eggs were directly deposited onto cryo-EM grids. Following a rapid, blotting-based rupture step directly on the EM-grid^16^, the egg cytoplasm formed a thin, electron-transparent film across the grid holes, followed by immediate vitrification within seconds (Fig.1a and Extended Data Fig. 2a). Raw cryo-EM micrographs of these preparations revealed a dense network of distinct filament-like structures, confirming the preservation of the CPL (Fig. 1b). Untilted, high-resolution micrographs were then collected for structure determination by the single particle cryo-EM approach.

**Fig. 1.**
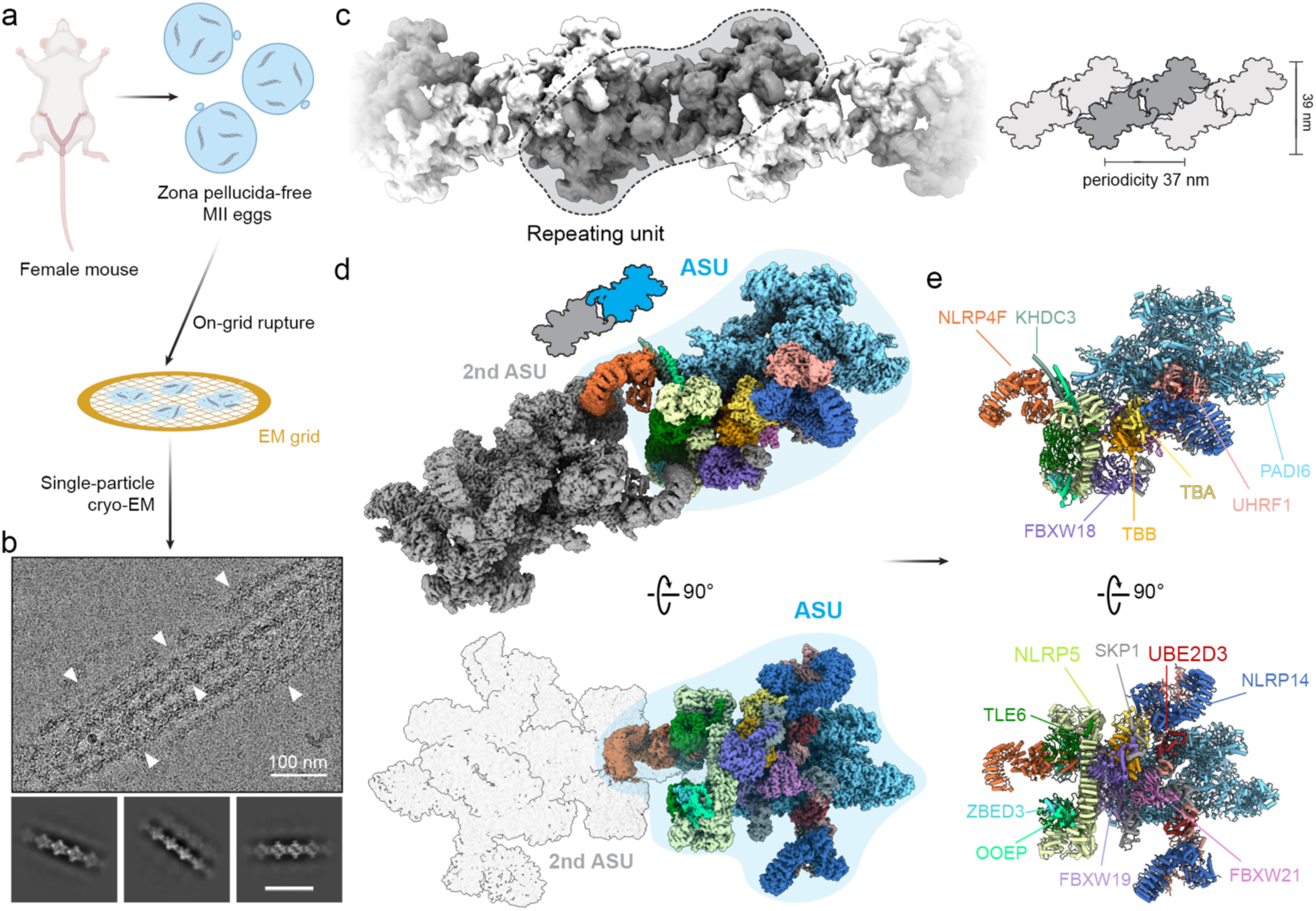
Native cryo-EM structure of the cytoplasmic lattice from mouse MII eggs. **a,** Schematic of the purification-free cryo-EM workflow. Zona pellucida-free MII eggs were applied directly to EM grids and mechanically ruptured on-grid to generate a thin layer of cytoplasmic content for single-particle cryo-EM analysis. **b,** Representative cryo-EM micrograph showing native CPL filaments. Arrowheads indicate individual filaments. Representative 2D class averages are shown below. Scale bar: 100 nm. **c,** Reconstruction of the CPL filament reveals an extended filamentous architecture with a periodicity of ∼37 nm. One repeating unit is outlined. Right, schematic representation of the periodic organization of the filament. **d,** Density map of the CPL repeating unit shown in two views. One ASU is highlighted in color. **e,** Atomic model of the CPL ASU shown in two orthogonal views, with components colored and labeled.

Initial 3D reconstruction demonstrated that the CPL adopts an extended filamentous organization composed of regularly repeating structural segments with an apparent C2 symmetry, with a periodicity of ∼37 nm (Fig. 1c). The repeating unit of the CPL filament can in principle be defined in two ways, based on two distinct local 2-fold symmetry axes along the filament (denoted symcenter 1 and 2 in Extended Data Fig. 3a). To identify the biologically relevant assembly unit, we performed 3D classification analysis centered on each symcenter separately. Classification on the assembly unit defined by symcenter 1 yielded a heterogeneous population, including a major class where half of the assembly was completely absent, indicating an unstable unit. By contrast, the symcenter 2-based assembly unit consistently yielded structurally complete classes, which differed only by modest conformational flexibility. This is consistent with contact-surface analysis, which confirmed that the symcenter-2 unit’s interface is double the size of the symcenter-1 unit. (Extended Data Fig. 3b). Consequently, we define this segment as the biological repeating unit of the CPL (Fig. 1c). Based on this definition, we further extracted the minimal asymmetric unit (ASU) of the C2 assembly unit and obtained a consensus reconstruction of the ASU at an overall resolution of 3.5 Å. Subsequent focused refinement of the functional units within the ASU further improved the map quality, yielding local resolutions up to 3.0 Å (Fig. 1d and Extended Data Fig. 2b).

The high-quality, native CPL cryo-EM maps enabled unambiguous assignment of the complete CPL protein inventory and accurate atomic model building (Fig. 1e). By integrating AlphaFold3 predictions^17^ and structure-similarity searches^18^ with the experimental density, we defined the complete molecular composition of the CPL (Fig. 2a and Table 1). The resulting model revealed a large assembly of 16 distinct protein components in specific copy numbers, with bound ligands and coordinated metals of functional implications. The structure establishes that the intricate macromolecular assembly of the MII CPL is organized into three functional modules: a scaffold of maternal-effect proteins (PADI6, TLE6, NLRP5, OOEP, KHDC3, NLRP4F, ZBED3, and NLRP14), ubiquitination machinery (a complete E3 ligase UHRF1 and E2 conjugating enzyme UBE2D3 module, and three FBXW-family E3 ligases, including FBXW18, FBXW19, and FBXW21, each paired with SKP1), and a tubulin assembly unit (αβ-tubulin heterodimer). To support our structural model, we performed quantitative proteomics analysis (Supplementary Table 1) and confirmed that all modeled CPL components were present in the proteomics data, most ranking among the top 10% highly expressed proteins in the egg proteome. These fundamental components form a minimal ASU, which undergoes dimerization and translational polymerization to generate a CPL filament. Multiple filaments crosslink into a higher-order fiber network (Fig. 2a).

**Fig. 2.**
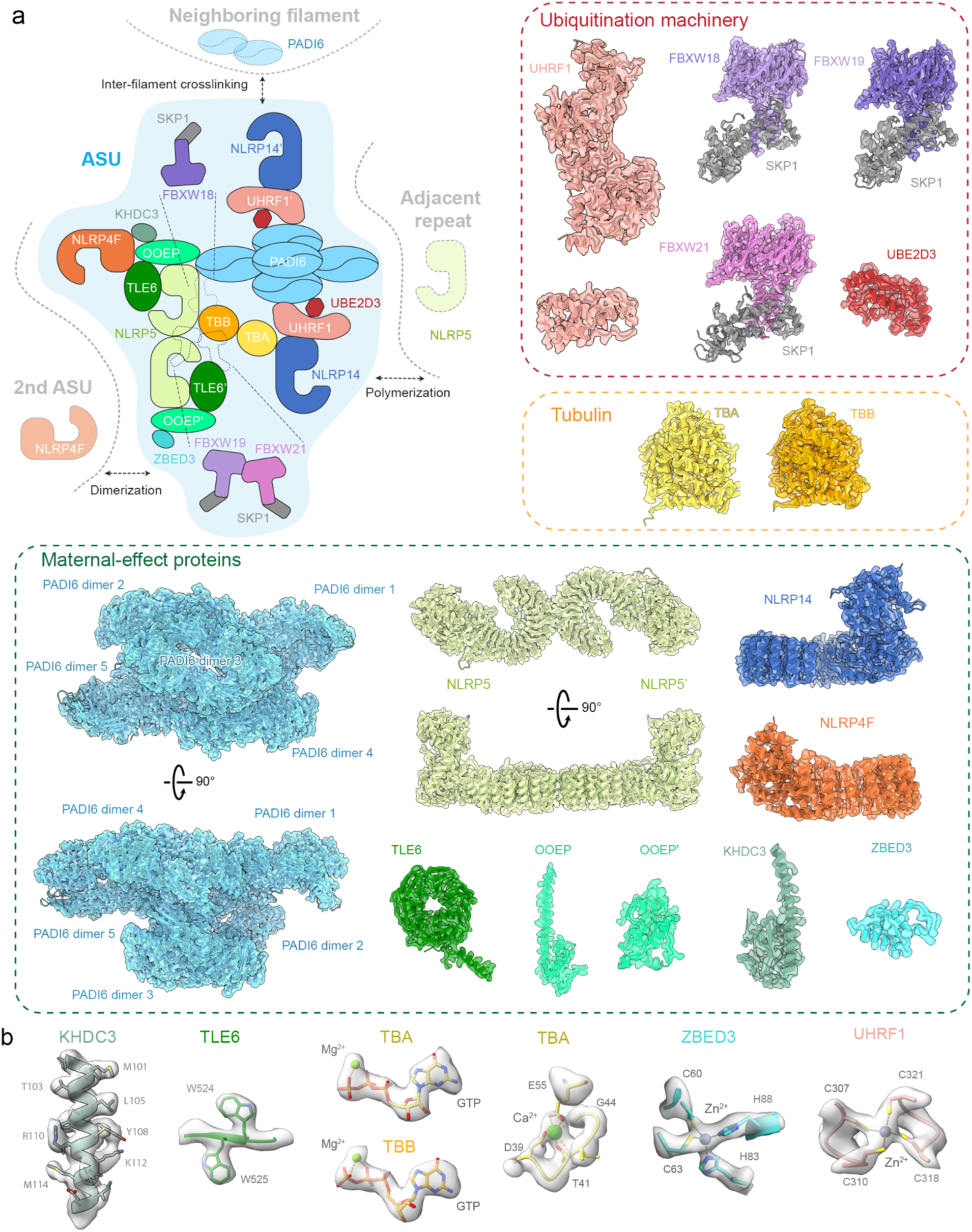
Architecture and functional modules of the CPL. **a,** Top left, schematic summary of the CPL ASU and its higher-order organization. Top right and bottom, atomic models of individual CPL components fitted into their corresponding cryo-EM map densities. The cryo-EM maps are displayed as transparent surfaces. The components are grouped by functional modules. Top right, ubiquitination machinery. Middle right, tubulin. Bottom, maternal-effect proteins. **b,** Representative high-resolution features of the native CPL cryo-EM map, including amino acid side chains, bound ligands and coordinated metal ions.

**Table 1.** Complete molecular inventory of the native CPL ASU determined by cryo-EM.

| Protein name | UNIPROT ID | Gene ID (MGI) | Number of residues | Number of copies per ASU<sup>a</sup> | Proteomics ranking | Protein category |
| --- | --- | --- | --- | --- | --- | --- |
| PADI6 | Q8K3V4 | 2655198 | 682 | 10 | 1 | Maternal protein |
| NLRP5 | Q9R1M5 | 1345193 | 1163 | 2 | 4 | Maternal protein |
| TLE6 | Q9WVB3 | 2149593 | 581 | 2 | 27 | Maternal protein |
| OOEP | Q9CWE6 | 1915218 | 164 | 2 | 118 | Maternal protein |
| KHDC3 | Q9CWU5 | 1914241 | 440 | 1 | 12 | Maternal protein |
| ZBED3 | Q9D0L1 | 1919364 | 228 | 1 | 46 | Maternal protein |
| NLRP4F | L7N1W9 | 2145528 | 937 | 1 | 28 | Maternal protein |
| NLRP14 | Q6B966 | 1924108 | 993 | 2 | 11 | Maternal protein |
| UHRF1 | Q8VDF2 | 1338889 | 782 | 2 | 14 | Ubiquitination |
| UBE2D3<sup>b</sup> | P61079 | 1913355 | 147 | 2 | 434 | Ubiquitination |
| FBXW18 | Q3TSA9 | 3505704 | 469 | 1 | 77 | Ubiquitination |
| FBXW19 | Q8C2W8 | 3505706 | 466 | 1 | 51 | Ubiquitination |
| FBXW21 | Q8BI38 | 2443323 | 468 | 0.6 | 84 | Ubiquitination |
| SKP1 | Q9WTX5 | 103575 | 163 | 2.6 | 41 | Ubiquitination |
| TBA1C<sup>c</sup> | P68373 | 1095409 | 449 | 1 | 22 | Tubulin |
| TBB4B<sup>c</sup> | Q9D6F9 | 107848 | 444 | 1 | 21 | Tubulin |

<sup>a</sup> Values represent the number of copies per asymmetric unit (ASU). <sup>b</sup> Modeled here as UBE2D3, although other highly similar UBE2D family members cannot be definitively excluded. <sup>c</sup> Specific tubulin isoforms could not be distinguished from the cryo-EM data. TBA1C and TBB4B are used as representatives for model building.

### Maternal-effect proteins form the scaffold of CPL

Maternal-effect proteins constitute the main structural framework of the CPL. This framework is organized into two linked modules: a central core of a dimer of NLRP5–TLE6–OOEP heterotrimer with additional associated components and a peripheral assembly of five PAID6 dimers. Together, these modules define the overall architecture of the CPL ASU and provide the scaffold for recruitment of the remaining functional CPL components.

At the center of the CPL, two sets of NLRP5, TLE6, and OOEP assemble into a heterodimeric module. This module is built around two NLRP5 molecules, which dimerize back-to-back through their leucine-rich repeat (LRR) domains (Extended Data Fig. 4a), consistent with a previously determined structure^19^. From a common NLRP5–TLE6–OOEP scaffold, the two halves diverge in their associated factors. In one half, the scaffold-associated NLRP4F engages KHDC3 through its N-terminal domain, whereas in the other, the corresponding NLRP4F binds ZBED3 through its C-terminal domain (Extended Data Fig. 4a).

The peripheral scaffold of the CPL is formed by a pentameric array of PADI6 homodimers (PADI6 dimer 1-5, Fig. 2a and Extended Data Fig. 4b), each adopting a head-to-tail configuration. This assembly is stabilized by two classes of inter-dimer interactions: contacts between the N-terminal Ig-like domains and packing between the C-terminal α/β-propeller domains (Extended Data Fig. 4b). Through its α/β-propeller domain, PADI6 dimer 1 contacts NLRP5 in the core scaffold, thereby linking the peripheral PADI6 assembly to the central NLRP5–TLE6–OOEP scaffold.

### The CPL sequesters specialized ubiquitination modules

One of the most prominent features of the CPL structure is its abundance of ubiquitin enzymes. Notably, the structure reveals two distinct classes of sequestered ubiquitination machinery. The first is a UHRF1–UBE2D3 module, comprising the E3 ligase UHRF1 and its cognate E2 enzyme UBE2D3 (Fig.3a). The E3 and E2 enzymes form a discrete module anchored between the scaffold NLRP14 and PADI6 dimer 2 in a CPL ASU. 3D classification further showed that in ∼61% of particles, another copy of UHRF1–UBE2D3–NLRP14 assembly is positioned beneath the CPL core through interaction with PADI6 dimer 4, forming an underfoot module (Extended Data Fig. 3c).

UHRF1 is an established epigenetic regulator in early development for histone ubiquitination and DNA methylation maintenance^20,21^. Structurally, UHRF1 is a multidomain protein composed of five major domains connected by flexible linkers. In the CPL-bound state, however, all five domains are well resolved and stably docked within the assembly (Fig. 3a). Its function normally depends on accessible chromatin-recognition and catalytic surfaces: the PHD domain binds the histone H3 tail, and the SRA domain recognizes hemi-methylated DNA. Inspection of the CPL-bound structure, however, shows that these surfaces are occluded by neighboring CPL components (Fig. 3b). This observation supports that the architecture of the CPL traps the UHRF1 module in a structurally constrained, inactive state.

**Figure 3.**
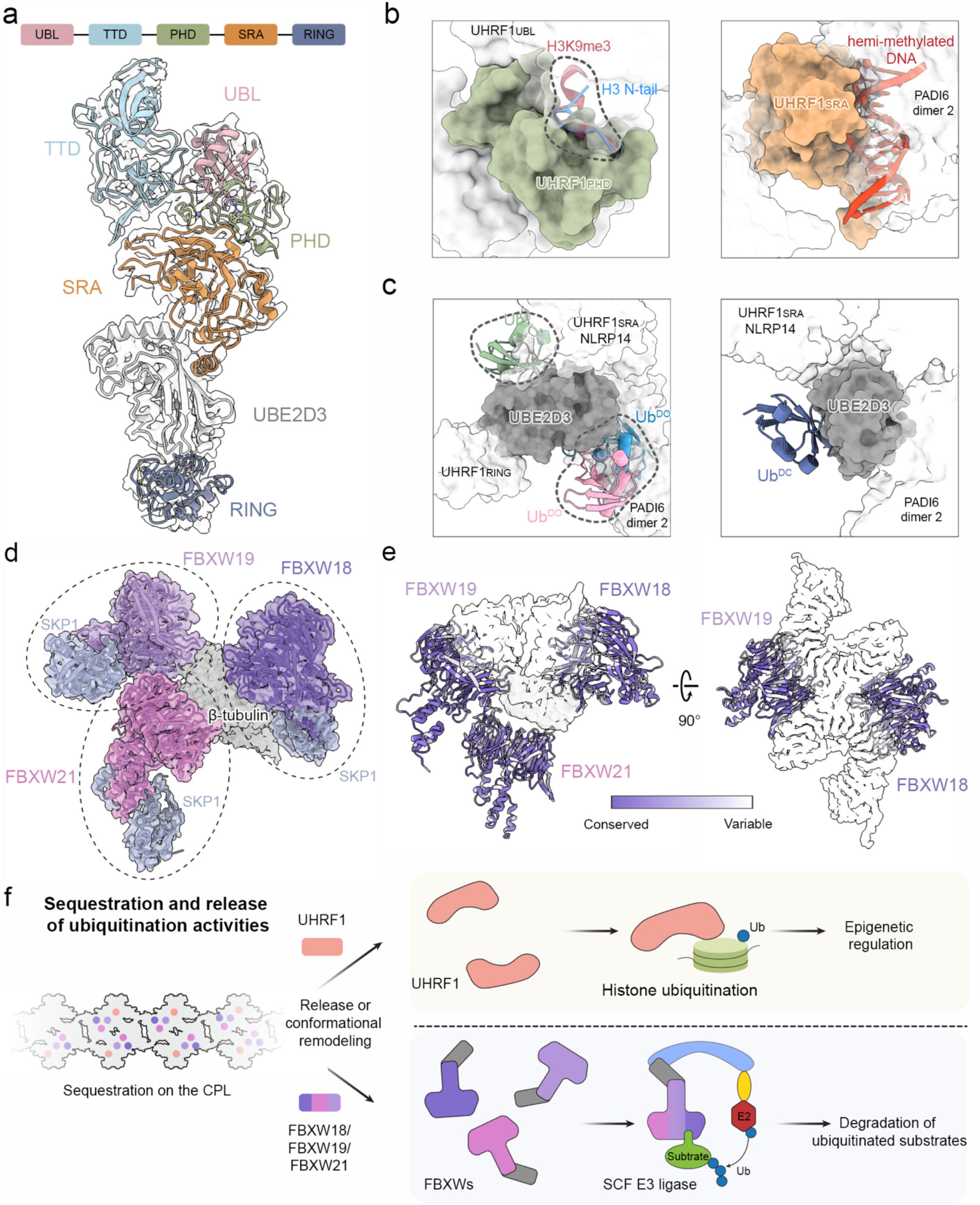
The CPL sequesters specialized ubiquitination machinery. **a,** Overall structure of the CPL-bound UHRF1–UBE2D3 E3-E2 module. **b**, Canonical chromatin-recognition surfaces of UHRF1 are occluded in the CPL context. Left, superposition of previously reported H3 N-tail- (PDB ID: 3SOU) and H3K9me3- (PDB ID: 4GY5) bound ligands onto the CPL-bound UHRF1 domain shows steric clash (highlighted by dashed contour) with surrounding CPL components. Right, superposition of hemi-methylated DNA onto the CPL-bound UHRF1 domain (PDB ID: 3CLZ) shows that the canonical DNA-binding mode is incompatible with the assembled CPL environment. **c**, Canonical ubiquitin-binding modes of UBE2D3 modeled in the CPL context. Left, backside-bound ubiquitin (Ub^B^, PDB ID: 4v3l) and donor ubiquitin in the open conformation (Ub^DO^, PDB IDs: 5ifr and 5d0k) are sterically occluded by surrounding CPL components (highlighted by dashed contours). Right, donor ubiquitin in the closed conformation can be accommodated (Ub^DC^, PDB ID: 4v3l). d, Three distinct FBXW–SKP1 modules (marked by ovals) are positioned at defined sites within the CPL ASU. FBXW18, FBXW19 and FBXW21 are shown in different shades of purple, and SKP1 is shown in grey. e, Mapping of sequence conservation onto the three CPL-associated FBXWs. Residues that vary among FBXW18, FBXW19 and FBXW21 cluster at interfaces with neighboring CPL components. f, Schematic model for sequestration and release of ubiquitination activities by the CPL.

UHRF1 makes extensive direct contact with UBE2D3, effectively cradling the E2 enzyme within the CPL assembly, which is consistent with the previously reported PADI6–UHRF1–UBE2D complex^22^. To assess whether the canonical ubiquitin-binding modes of UBE2D3 can be accommodated in this context, we performed structural modeling. This analysis showed that both the backside-bound ubiquitin (Ub^B^)^23^ and the donor ubiquitin in the open conformation (Ub^DO^)^24,25^ would sterically clash with surrounding CPL components, whereas donor ubiquitin in the closed conformation (Ub^DC^) can be accommodated (Fig. 3c). Thus, the architecture of the CPL appears to restrict UBE2D3 to a catalytically-poised E2∼Ub state, while disfavoring more accessible ubiquitin-binding configurations.

A second major class of ubiquitin ligases sequestered within the CPL comprises three distinct FBXW-family proteins, each paired with one SKP1 (Fig. 3d). The FBXW proteins act as substrate receptors, and SKP1 links them to the core ligase scaffold in canonical SCF (SKP1-CUL1-F-box) E3 ligase complexes, which govern turnovers of essential proteins during embryonic development^26,27^. Two stably incorporated FBXW–SKP1 assemblies contact two scaffold NLRP5 molecules together with one β-tubulin, and focused classification showed that a third FBXW is present in ∼60% of CPL units (Extended Data Fig. 3d). All three contain the characteristic eight-bladed WD40 domain and are highly similar to one another (Extended Data Fig. 5a). Although homologous FBXW-family members could be fitted into these densities with comparable overall agreement, and multiple leading candidates are all highly expressed in the oocytes and early embryos (Extended Data Fig. 5b and c), the well-resolved side-chain features in our cryo-EM map allowed confident assignment of three members, FBXW18, FBXW19, and FBXW21 (Extended Data Fig. 5d).

To understand why these three FBXW-family members are incorporated into the CPL, we mapped sequence variation onto the structure. The variable residues cluster at interfaces with neighboring CPL components (Fig. 3e), indicating that family-specific sequence features help determine CPL binding. This supports selective recruitment of a specialized FBXW subset rather than nonspecific capture of closely related homologs. Structural comparisons with well-characterized FBXW homologs indicate that the conserved substrate-binding site lies on the top surface of the WD40 β-propeller (Extended Data Fig. 6a). In the CPL-bound state, this canonical substrate-binding pocket is buried within the assembly and inaccessible (Extended Data Fig. 6b).

These observations indicate that CPL selectively sequesters a highly specialized set of ubiquitination factors, thereby preventing premature ubiquitination in the egg while preserving these activities for later developmental use. The captured machinery points to two distinct functional outputs: UHRF1-mediated chromatin regulation and FBXW-containing, SCF-dependent substrate degradation. Together with the specificity of FBXW molecules bound to the CPL, this suggests that the CPL is not a generic storage depot, but a dormant and highly specific ubiquitination platform poised to orchestrate protein homeostasis and chromatin remodeling during early embryonic development (Fig. 3f).

### The CPL functions as a reservoir for assembly-competent tubulin

In addition to serving as a reservoir for ubiquitin enzymes, our CPL structure suggests another potentially important storage function: sequestration of tubulin, the fundamental building block of microtubules^28^. We observed two globular densities that could be unambiguously assigned as an αβ-tubulin heterodimer (Fig. 4a), positioned between the maternal-effect proteins NLRP14 and NLRP5. Inspection of the nucleotide-binding pockets showed that both the α- and β-subunits are bound to guanosine triphosphate (GTP) together with a coordinating magnesium ion (Fig. 4a). Whereas α-tubulin constitutively binds GTP, the GTP bound to β-tubulin is hydrolyzed to GDP upon incorporation into the microtubule lattice^29,30^. The CPL-bound heterodimer is therefore captured in a pre-hydrolysis state before microtubule assembly.

**Figure 4.**
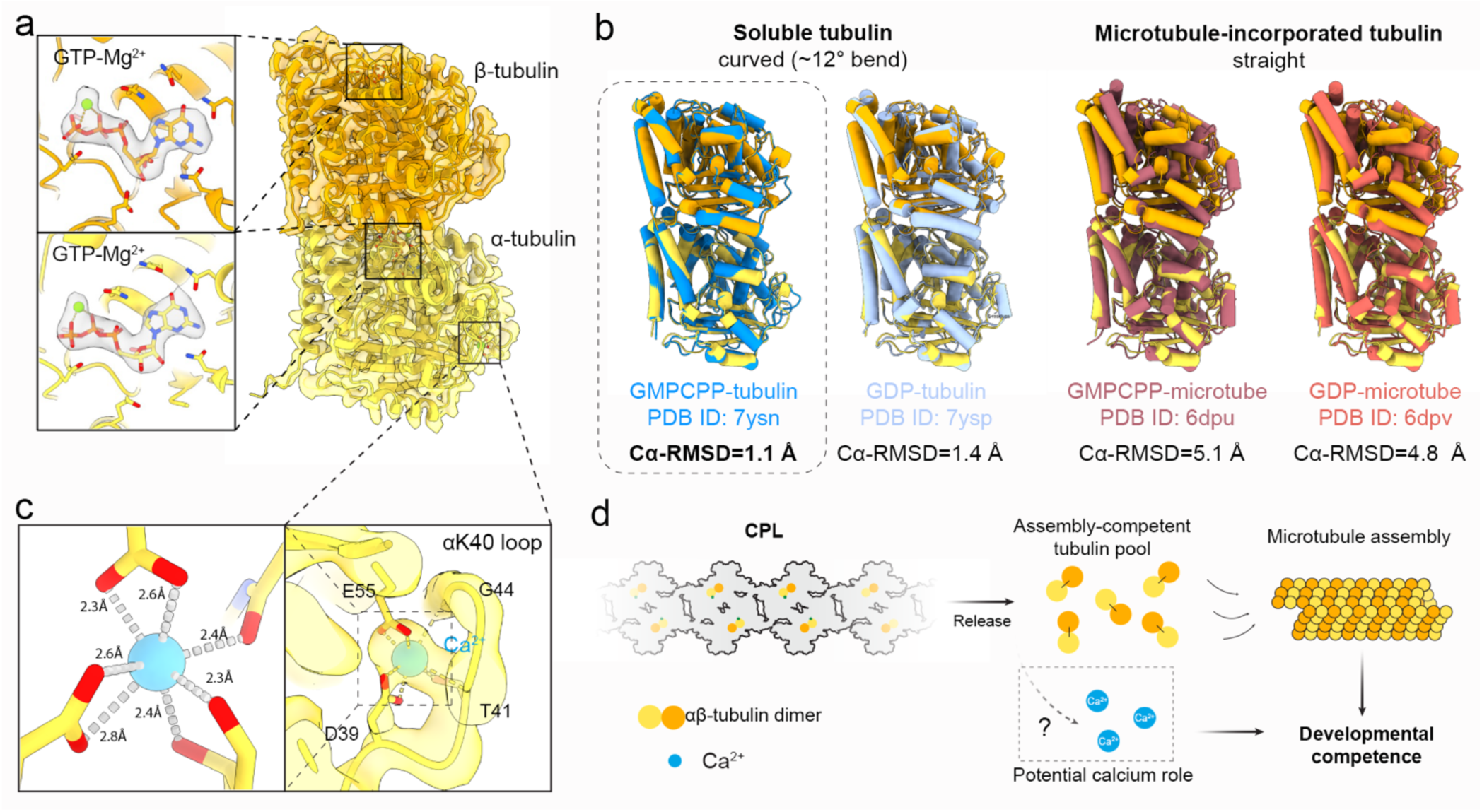
CPL stores assembly-competent αβ-tubulin. **a,** Structure of the CPL-associated αβ-tubulin heterodimer. Insets show map densities for GTP and Mg²⁺ bound to both the α- and the β-subunits. **b,** Structural comparison of CPL-bound tubulin with representative tubulin conformations in distinct nucleotide and assembly states. GMPCPP is a non-hydrolysable GTP analogue. After superposition of α-tubulin, the Cα-RMSD of β-tubulin was calculated. **c**, A coordinated Ca²⁺ ion bound by the α-tubulin K40 loop. Insets show the coordination environment and coordination geometry of the Ca²⁺ ion. **d,** Schematic model for CPL-mediated sequestration of tubulin to support developmental competence.

To further define the structural state of CPL-bound tubulin, we compared the sequestered αβ-tubulin heterodimer with tubulin in soluble and microtubule-assembled states (Fig. 4b). Soluble αβ-tubulin is characteristically curved, with an approximately 12° bend, whereas incorporation into the microtubule lattice straightens the heterodimer^31,32^. Using α-tubulin for alignment, we measured the Cα root mean square deviation (Cα-RMSD) of β-tubulin as a readout of heterodimer conformation. The CPL-bound heterodimer matched most closely to the curved soluble state, especially GTP analogue-bound tubulin, and differed markedly from the straight conformation found in the microtubule lattice (Fig. 4b). Together with its nucleotide state, this shows that the CPL captures tubulin in a pre-hydrolysis conformation characteristic of soluble, assembly-ready αβ-tubulin. Quantitative proteomics further showed that a substantial fraction of total cellular tubulin, in the range of 12-14%, is docked on the CPL (Supplementary Note 1). These findings support a model in which the CPL functions as a tubulin reservoir, sequestering the heterodimers in their assembly-competent state. Such sequestration is expected to prevent spontaneous microtubule polymerization driven by high cytosolic tubulin concentrations, while preserving a reserve that can be rapidly deployed for cytoskeletal remodeling after fertilization (Fig. 4d).

Notably, our structure also uncovered a well-defined density corresponding to a metal ion coordinated in the α-tubulin Lys40 (αK40) loop (Fig. 4c). The metal ion is positioned in an electronegative pocket formed by residues D39, T41, G44, and E55 of α-tubulin, with coordination geometry and ligand distances most consistent with calcium^33^. A similar Ca²⁺ state coordinated by the αK40 loop (Extended Data Fig.7) was reported in multiple tubulin structures in previous studies^34–36^, but was suggested as crystallization artifact^37^. Our native structure lends support to the potential physiological relevance of this Ca²⁺-bound tubulin state. In microtubules, the αK40 loop is a conserved regulatory element containing the canonical K40^38^, and it is highly flexible and poorly resolved^37^. The well-defined Ca²⁺-coordinating conformation observed here may be favored in the CPL-sequestered state. Although the physiological significance remains to be established, these observations raise the possibility that CPL-associated tubulin may contribute to a calcium-related role for egg activation at fertilization^41^.

### Higher-order assembly of CPL fibers

Given the CPL’s likely role as a major reservoir for ubiquitin enzymes and tubulin, it is important to understand how CPL filaments assemble into higher-order fibers capable of accommodating the many copies of these components. To define the assembly mechanism, we docked the repeating-C2 unit into a cryo-EM map of the extended filament using rigid-body fitting. Adjacent repeating units stack in a strictly coaxial manner (Fig. 5a, b and Extended Data Fig. 8a), generating a linear filament without intrinsic curvature or helical twist. This longitudinal linkage is mediated by relatively rigid contacts between PADI6 dimer 5 of one repeating unit and an NLRP5 module of the next (Fig. 5a). In contrast to the rigidity of the inter-unit interface, flexibility resides within each repeating unit. The two ASUs within the C2 unit can move relative to one another. Alignment of one ASU showed that the other undergoes continuous swinging motions in three dimensions, with the two NLRP4F molecules acting as hinges (Fig.5b and Extended Data Fig. 8b). These local motions could propagate to generate curved CPL filaments observed in the TEM micrographs (Extended Data Fig. 1a).

**Figure 5.**
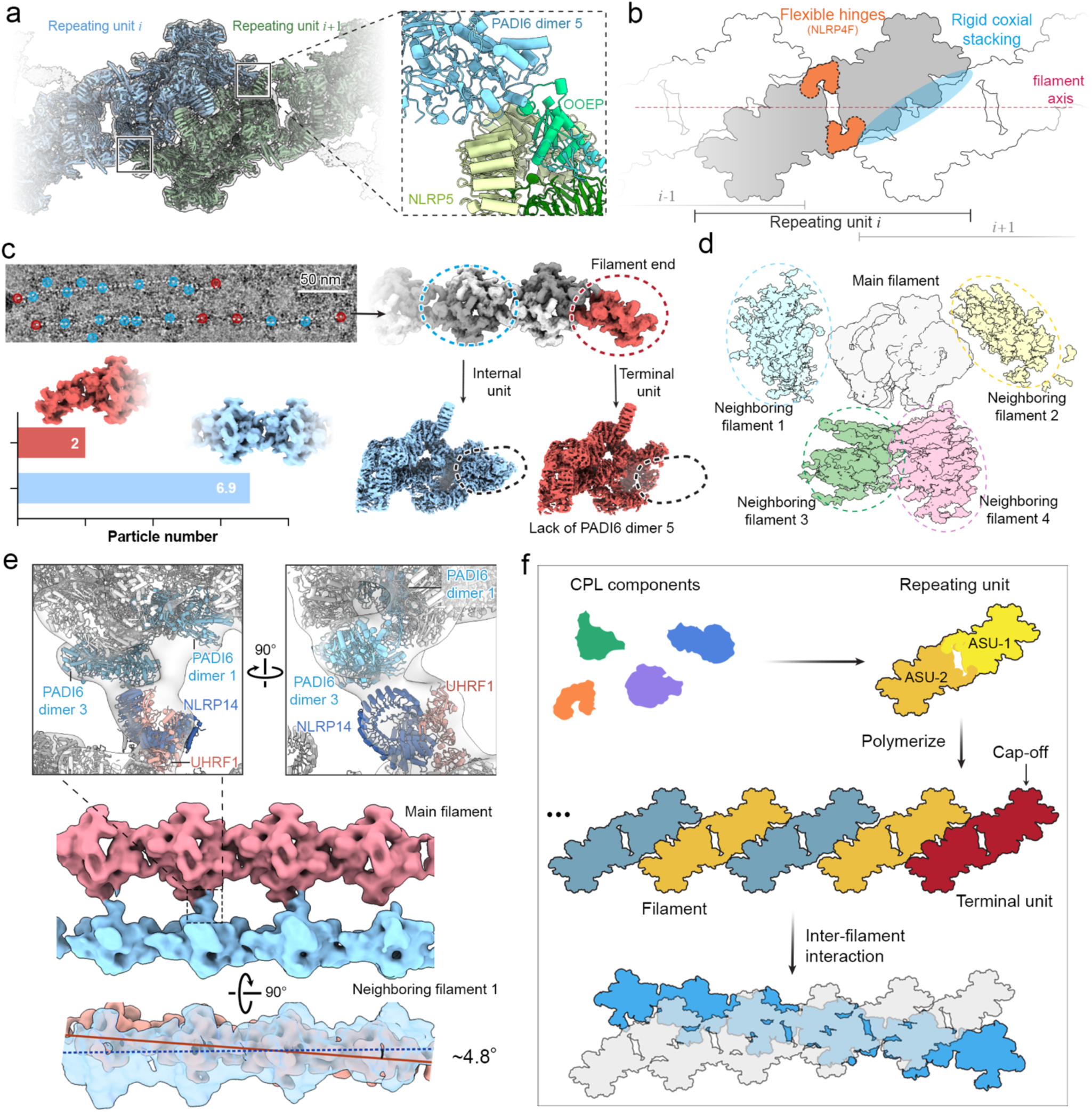
Structural basis of CPL filament assembly, termination, and higher-order network formation. **a,** Cryo-EM map showing the longitudinal polymerization of adjacent repeating units (*i* and *i*+1). The zoomed-in atomic model reveals that this coaxial stacking is primarily driven by specific interactions between the PADI6 dimer 5 of one unit and the NLRP5 of the adjacent unit. **b,** Schematic representation of the organization of the CPL filament and its structural flexibility. **c**, Structural analysis reveals how filaments end and are capped by a terminal unit. Top left: a representative micrograph shows the location of internal (blue) and terminal (red) units. Bottom left: particle ratio of internal and terminal units. Right: 3D reconstruction of the terminal unit (red, right) reveals the absence of PADI6 dimer 5 (marked by ovals). **d**, Cross-sectional view of the higher-order CPL 3D lattice reconstruction, demonstrating that a central main filament can potentially coordinate with up to four adjacent filaments positioned at its bottom and lateral sides. **d,** Cryo-EM maps and atomic models of filament pair reveal that inter-filament interaction is mediated by contacts between PADI6 molecules (dimers 1 and 3) of the main filament and the NLRP14-UHRF1 module of the adjacent filament. A side view of the reconstructed filament pair (bottom) demonstrates a relative crossing angle of ∼4.8° between the two filaments. **f**. A schematic model summarizing the hierarchical assembly of the 3D CPL network.

We next investigated how filament growth terminates. Classification analysis demonstrated that the filament terminates in a single ASU (half of a C2 repeating unit) (Fig. 5c), further supporting our definition of the ASU as the fundamental building block. Mapping these half-units back to the micrographs showed that they localize predominantly to the filament ends. The ratio of the half-terminal units to the intact internal units was ∼2:7, suggesting that an average filament contains about eight repeating units. Given the ∼37 nm axial periodicity, this corresponds to an average filament length of ∼333 nm, consistent with the length measurements observed by TEM (Extended Data Fig. 1b). Reconstruction of the terminal unit further showed that it lacks PADI6 dimer 5, the structural element required for longitudinal stacking, indicating that its absence acts as an intrinsic stop to terminate polymerization (Fig. 5c).

Extended filament reconstructions additionally revealed connecting densities from neighboring filaments around the central filament, suggesting that one filament can engage up to four adjacent filaments at lateral and basal positions (Fig. 5d). Reconstruction of an interacting filament pair showed that this higher-order connection is mediated mainly by the basal NLRP14–UHRF1 module of one filament and lateral PADI6 dimers 1 and 3 of the neighboring filament (Fig. 5e, top and middle). The two filaments intersect at a shallow angle of ∼4.8° (Fig. 5e, bottom), an arrangement that may favor three-dimensional network formation.

Together, these data support a hierarchical model for CPL assembly (Fig. 5f): coaxial stacking of repeating and terminal units generates an apolar, end-capped filament, and directional lateral contacts crosslink these filaments into higher-order fibers, ultimately building the large, three-dimensional CPL network essential for early embryonic development.

## Discussion

The high-resolution, native structures of the mouse CPL we determined in unfertilized eggs reveal the complete molecular composition, the assembly, and architectural principles of the lattice. The functional capacity of the CPL is directly suggested by our structures.

The CPL is an apparently dormant ubiquitination hub in eggs, incorporating four inactivated E3 ubiquitin ligases and one E2 ubiquitin conjugating enzyme. UHRF1 is responsible for ubiquitination of histone H3 to maintain DNA methylation during early embryogenesis. In the UHRF1–UBE2D E3-E2 module, the substrate- and ubiquitin-binding pockets are mostly sterically occluded by the surrounding subunits of the CPL. Anchoring this module to the CPL restricts UHRF1 in the cytoplasm, providing spatial and temporal regulation of UHRF1 for its nuclear translocation and epigenetic reprogramming. The oocyte-specific FBXW-family E3 ubiquitin ligases, FBXW18, FBXW19, and FBXW21, are bound to the CPL at three distinct positions, each paired with one adaptor protein SKP1. Their putative substrate-binding pockets are buried within the CPL, rendering their activities inaccessible in eggs. Although the substrates for these E3 ligases remain elusive, we speculate that structural remodeling of the CPL or the release of these E3 ligases after fertilization may facilitate targeted, ubiquitin-mediated degradation of paternal and/or maternal proteins during egg-to-embryo transition.

The CPL may serve as a reservoir of tubulin for cytoskeletal remodeling. The αβ-tubulin heterodimers anchored in the CPL are bound to GTP-Mg^2+^, adopting the curved, pre-hydrolysis conformation characteristic of soluble, assembly-ready tubulin and distinct from that in polymerized microtubules. Remarkably, the CPL sequesters ∼12-14% of the total tubulin in eggs in a non-polymeric but readily deployable state to prevent spontaneous microtubule assembly. We propose that the CPL serves as a reserve for microtubule polymerization, supporting the rapid, *de novo* assembly of an extensive cytoplasmic microtubule network that drives early pronuclear migration in mammalian zygotes^42,43^.

Across the many cryo-EM structures of free tubulin and microtubule polymers reported to date, the α-tubulin K40 loop has remained disordered. It is therefore notable that, in the CPL, we observe a stable K40 loop coordinated with a putative calcium ion. Given the abundance of CPL-bound α-tubulin, the total CPL-associated calcium could, in principle, be comparable to the fertilization-induced calcium spikes. This raises the prospect that the CPL contributes to calcium regulation during fertilization and embryonic development, a possibility that will be important to test in future studies.

The translationally stacked architecture of the CPL appears suitable for cargo sequestration and autoinhibition, resembling the assembly principles of several metabolic enzyme filaments^44,45^. This organization may be particularly suited to the extraordinary lifespan of mammalian oocytes. In many species, the oocytes are formed before birth and maintained in prolonged arrest^46^, during which CPLs must preserve their cargo for eventual fertilization. Canonical cytoskeletal polymers such as actin filaments, microtubules, and ESCRT-III are typically linked to regulated remodeling^47^. In this context, the physiological cues and structural transitions that govern how CPL components are activated and deployed during fertilization and early embryogenesis, or when and how CPLs disassemble remain unknown. Whether disassembly occurs rapidly or progressively, and whether and how cargo deployment is coordinated with specific stages of early embryogenesis, remain to be determined. The new experimental methodology described here, together with established quantitative cell biology approaches for studying pre-implantation embryonic development, will help address these open questions.

Our approach of directly depositing individual eggs onto cryo-EM grids and rupturing them to expose cytoplasmic materials, while preserving native organization opens new avenues to study the ultrastructural basis of cellular architecture in large, long-lived cells. Given that non-synonymous mutations in CPL genes are linked to female infertility, this strategy provides a foundation for single-cell structural analysis of human eggs and embryos at high resolution, with potential to illuminate currently inaccessible aspects of human reproduction and inform assisted reproductive technologies.

## Methods

### Mouse husbandry, oocyte collection, and *in vitro* maturation into metaphase II eggs

Fully grown, prophase I–arrested oocytes were collected from the ovaries of 6-8-week-old CD1 females in M2 medium supplemented with dibutyryl cyclic AMP (dbcAMP) to maintain meiotic arrest. Oocytes were released from arrest by washing through multiple droplets of dbcAMP-free M2 medium and matured *in vitro* into MII eggs as described previously^48^. Mature eggs were prepared for subsequent experiments by removing the ZP through sequential washing in droplets of Tyrode’s solution. Mice were maintained under specific pathogen-free conditions on a 12-hour light/dark cycle. All animal procedures were approved by the Yale University Institutional Animal Care and Use Committee (protocols 2024-20408 and 2024-20562).

### Negative Stain EM imaging

10-12 ZP-free eggs were transferred onto poly-L-lysine-coated culture dishes and allowed to settle and attach in PBS. Samples were then fixed in 2.5% Glutaraldehyde in 0.1 M sodium cacodylate buffer for 1 hour at room temperature. Samples were rinsed three times with 0.1 M sodium cacodylate and post-fixed in 1% osmium tetroxide with 0.8% potassium ferrocyanide in 0.1 M sodium cacodylate for 1 hour. Samples were then rinsed once in buffer and three times in water, then transferred to 2% uranyl acetate for 1 hour, followed by three rinses in distilled water. Dehydration was performed through a graded ethanol series (50%, 70%, 90%, and 100%), and samples were embedded in Embed-812 epoxy resin (EMS, PA). Ultrathin sections (60 nm) were cut using a Leica UC7 ultramicrotome. Grids were post-stained with 2% uranyl acetate and lead citrate before imaging on an 80 kV FEI Tecnai Biotwin TEM equipped with an AMT NanoSprint 15 camera.

### Tandem mass tag (TMT)-based quantitative proteomics of mouse eggs

For egg proteomics, 500 fully grown, prophase I-arrested oocytes were isolated from 8 C57BL/6 female mice aged 6-12 weeks. Oocytes were maintained in prophase arrest in M2 medium supplemented with dbcAMP, then released and matured *in vitro* to MII-arrested eggs as described previously^48^. Eggs were then washed three times through several droplets of PBS to remove residual medium and extracellular contaminants and lysed in the RIPA buffer. Protein extracts were processed for TMT-based quantitative proteomics. Peptides were generated by tryptic digestion, labeled with isobaric TMT reagents, and fractionated before liquid chromatography-tandem mass spectrometry (LC-MS/MS) analysis. Mass spectrometry data were acquired using high-resolution Orbitrap instrumentation, and raw data were processed using established pipelines as described previously^48^. Peptide and protein identifications were filtered to a false discovery rate of 1% using a target–decoy approach. Relative protein quantification was based on TMT reporter ion intensities, with normalization across channels to correct for labeling efficiency and sample loading. All proteomic sample preparation, TMT labeling, LC-MS/MS acquisition, and initial data processing were performed by the Proteomics Core Facility at the University of Bristol.

### Native cryo-EM sample preparation

R2/1 holey carbon gold grids (200 mesh, Quantifoil) coated with a 2-nm continuous carbon film were glow-discharged for 25 s at 20 mA. 5–7 ZP-free MII eggs were manually placed onto each grid. The grids were then processed using a Vitrobot Mark IV (Thermo Fisher Scientific) at 10 °C and 100% humidity. To achieve optimal ice thickness and egg rupture on the grid, a consecutive triple-blotting was employed with a blot force of 20 and sequential blot times of 1 s, 1 s, and 3 s. Following the final blot, the grid was immediately plunged and vitrified in liquid ethane.

### Cryo-EM data acquisition

Cryo-EM data were acquired on a Titan Krios G2 transmission electron microscope (Thermo Fisher Scientific) operated at 300 kV. The microscope was equipped with a Gatan BioQuantum energy filter (20-eV slit width) and a Gatan K3 direct electron detector. Automated data collection was performed using SerialEM software using the single particle cryo-EM approach. Micrographs were recorded in counting mode at a magnification of 81,000×, corresponding to a calibrated physical pixel size of 1.068 Å. Each movie was collected with a total dose of 50 e⁻/Å². The defocus range was set between −1.2 μm and −2.5 μm.

### Cryo-EM data processing

The acquired movies were processed for motion correction^49^ and contrast transfer function (CTF) correction^50^ using CryoSPARC^51^. A subset of 12,231 micrographs was first selected for particle picking. An initial particle set was generated by manual picking followed by 2D classification, and the resulting class averages were used for template-based automatic particle picking in CryoSPARC. After 2D classification, a clean dataset of 62,986 particles was retained. These particles were subjected to *ab initio* reconstruction and heterogeneous refinement. The most well-defined class was subjected to homogeneous refinement with C2 symmetry applied, resulting in a 4.8 Å resolution map that served as the reference for subsequent comprehensive particle picking.

High-resolution 2D template matching was performed on the entire dataset of 44,044 micrographs using GisSPA^52^, utilizing reference templates generated with a 3° angular step, bin2 scaling, and C2 symmetry. This process yielded an initial set of 491,871 particles. After the removal of duplicate particles, several rounds of local refinement and 3D classification, and subset curation, a clean dataset of 290,887 particles was obtained. These particles were recentered to symmetry center 2. Subsequent local refinement and CTF refinement yielded a repeating C2 unit map at an overall resolution of 4.1 Å. To obtain the ASU map, particles were then symmetry-expanded, recentered to the ASU center, and duplicate particles were removed. The resulting particles were refined in C1 symmetry using an ASU mask together with CTF refinement, producing an ASU consensus map at 3.5 Å resolution.

To improve local map quality and resolve heterogeneous regions, focused local refinement and focused 3D classification were performed with region-specific masks. Local refinement yielded maps for the NLRP14+UHRF1 (the UBL, TTD, PHD, and SRA domains) region at 3.1 Å resolution, the tubulin dimer at 3.1 Å resolution, PADI6 dimers 1–4 at 3.3 Å resolution, the NLRP5+TLE6+OOEP+ZBED3 region at 3.0 Å resolution, the NLRP5+TLE6+OOEP+KHDC3 region at 3.1 Å resolution, the NLRP4F region at 3.6 Å resolution, and the UBE2D3+UHRF1(RING) region at 3.6 Å resolution. Focused classification further resolved heterogeneity in several regions: PADI6 dimer 5 was present in 79% of particles and was refined to 3.5 Å resolution; the FBXW19+FBXW21 region separated into classes containing FBXW19 alone (42%) or both FBXW19 and FBXW21 (58%), with the latter refined to 3.3 Å resolution; the NLRP14′+UHRF1′+UBE2D3′ module was present in 61% of particles and refined to 4.2 Å resolution. The ASU composite map was generated by combining 10 locally refined maps. Application of C2 symmetry to the ASU composite map produced the final repeating unit composite map.

To analyze the extended filament structure, particles were re-extracted with a larger box size (2,048 pixels, bin4). Local refinement using a main-filament mask yielded an 8.6 Å resolution map of the filament. Subsequent 3D classification with an adjacent filament mask was performed to resolve the map of an interacting filament pair. The overall resolutions of all reconstructed maps were assessed using the gold-standard criterion of Fourier shell correlation^53^ at 0.143 cut-off^54^.

### Model building and refinement

Protein identification and model building were performed using a combination of template fitting, structure prediction, and structure-similarity searches. For the NLRP5–TLE6–OOEP core, the published structure (PDB ID: 8H93) was used as the initial model and docked into the map in ChimeraX^55^. For unassigned densities, candidate identities were evaluated in two ways. First, AlphaFold3-predicted models^56^ of candidate proteins were docked into the cryo-EM density and assessed for overall agreement. Second, when no clear candidate could be assigned a priori, the density was initially traced as a polyalanine model in Coot^57^. This polyalanine model was then used for a structure-similarity search against the AlphaFold database^56^ and the DALI server^18^ to identify structurally related candidates. Candidate subunits were subsequently fitted into the cryo-EM map, assembled into a composite model, and manually adjusted. Final model building was carried out iteratively in Coot, followed by real-space refinement in Phenix^58^ with secondary-structure, Ramachandran, and rotamer restraints to maintain model geometry and improve agreement with the map. The refinement statistics are in Extended Data Tables 1 and 2.

### Estimation of CPL-associated tubulin fraction

Protein abundances were obtained from TMT-based quantitative proteomics as summed reporter ion intensities (“area”). CPL-associated tubulin was estimated based on the structural stoichiometry, assuming a 1:1 relationship between tubulin heterodimers and the CPL components NLRP5, TLE6, and OOEP. Total tubulin abundance was calculated by summing all detected α- and β-tubulin isoforms. The fraction of CPL-associated tubulin was then calculated by normalizing CPL-derived tubulin estimates to total tubulin abundance. Detailed calculations are provided in Supplementary Note 1.

## Data availability

The cryo-EM density maps generated in this study have been deposited in the Electron Microscopy Data Bank under accession numbers EMD-76333 (CPL filament map), EMD-76335 (ASU consensus map), EMD-76334 (ASU composite map), EMD-76315 (focused on PADI6 dimers 1–4), EMD-76321 (focused on NLRP5+TLE6+OOEP+KHDC3), EMD-76322 (focused on NLRP5+TLE6+OOEP+ ZBED3), EMD-76323 (focused on NLRP14+UHRF1), EMD-76324 (focused on tubulin), EMD-76325 (focused on NLRP4F), EMD-76326 (focused on FBXW19+FBXW21), EMD-76327 (focused on UBE2D3+UHRF1 RING), EMD-76330 (focused on PADI6 dimer 5), and EMD-76331 (focused on NLRP14′+UHRF1′+UBE2D3′). The corresponding atomic model of CPL ASU has been deposited in the Protein Data Bank under accession number PDB ID: 12DL. The proteomics data generated in this study have been deposited in the PRIDE repository under accession number PXD076290. Source data and other materials are available from the authors upon reasonable request. Source data and other materials are available from the authors upon reasonable request.

## Acknowledgements

We thank Yale CryoEM Resource facilities for support with cryo-EM data collection. We thank the Yale Center for Cellular and Molecular Imaging (CCMI) EM facilities for assistance with negative-stain TEM of eggs. We thank Alice Sherrard at Yale University for helping with sample preparation for negative-stain TEM imaging. This work was supported in part by a Vallee Scholars Award (VS-2024-56), Pew Scholars Award (00037689) and a National Institutes of Health (NIH) grant (R35GM146725) to BM, a Lalor Foundation Postdoctoral Fellowship to JL, an NIH NICHD grant R00HD104924 and a David Sokal Innovation Award of Male Contraceptive Initiative 2024-303 to ST, and a Yale discretionary fund to YX.

## Author contributions

Conceptualization: BM, YX. Methodology: YL, JL, WZ, CW, ST, BM, YX. Investigation: YL, JL, WZ, CW. Visualization: YL, BM, YX. Project administration: BM, YX. Supervision: BM, YX. Funding acquisition: ST, BM, YX. Writing – original draft: YL, ST, BM, YX. Writing – review & editing: YL, ST, BM, YX.

## Open Access

This manuscript is the result of funding in part by the NIH and is subject to the NIH Public Access Policy. Through acceptance of this federal funding, NIH has been given the right to make this manuscript publicly available in PubMed Central upon the Official Date of Publication, as defined by NIH.

---

**Extended Data Figure 1.**
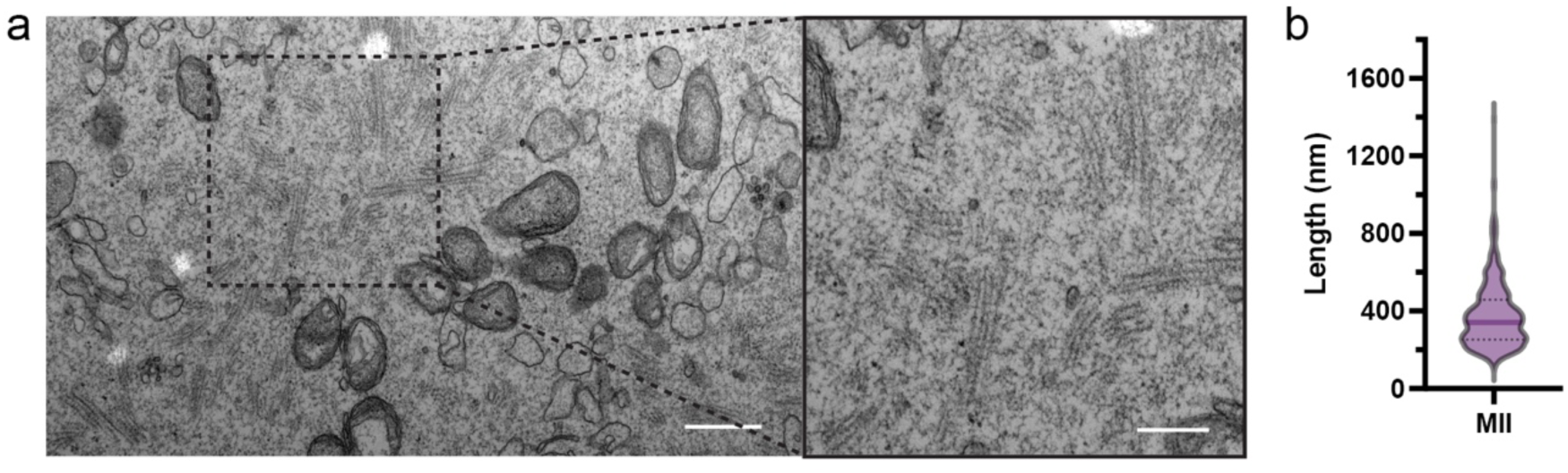
Macroscopic organization and quantification of the CPL network in mouse eggs. a, Representative TEM micrographs of ultrathin sections from intact mouse eggs. The right panel is a magnified view of the boxed region in the left panel. Scale bars, 600 nm (left) and 300 nm (right). **b**, Quantification of the lengths of individual CPL filaments measured from the TEM micrographs. The violin plot shows the distribution of filament lengths in eggs. The solid line indicates the median, and the width of the violin represents the relative density of measurements at each length. Length measurements were obtained from 465 filaments.

**Extended Data Figure 2.**
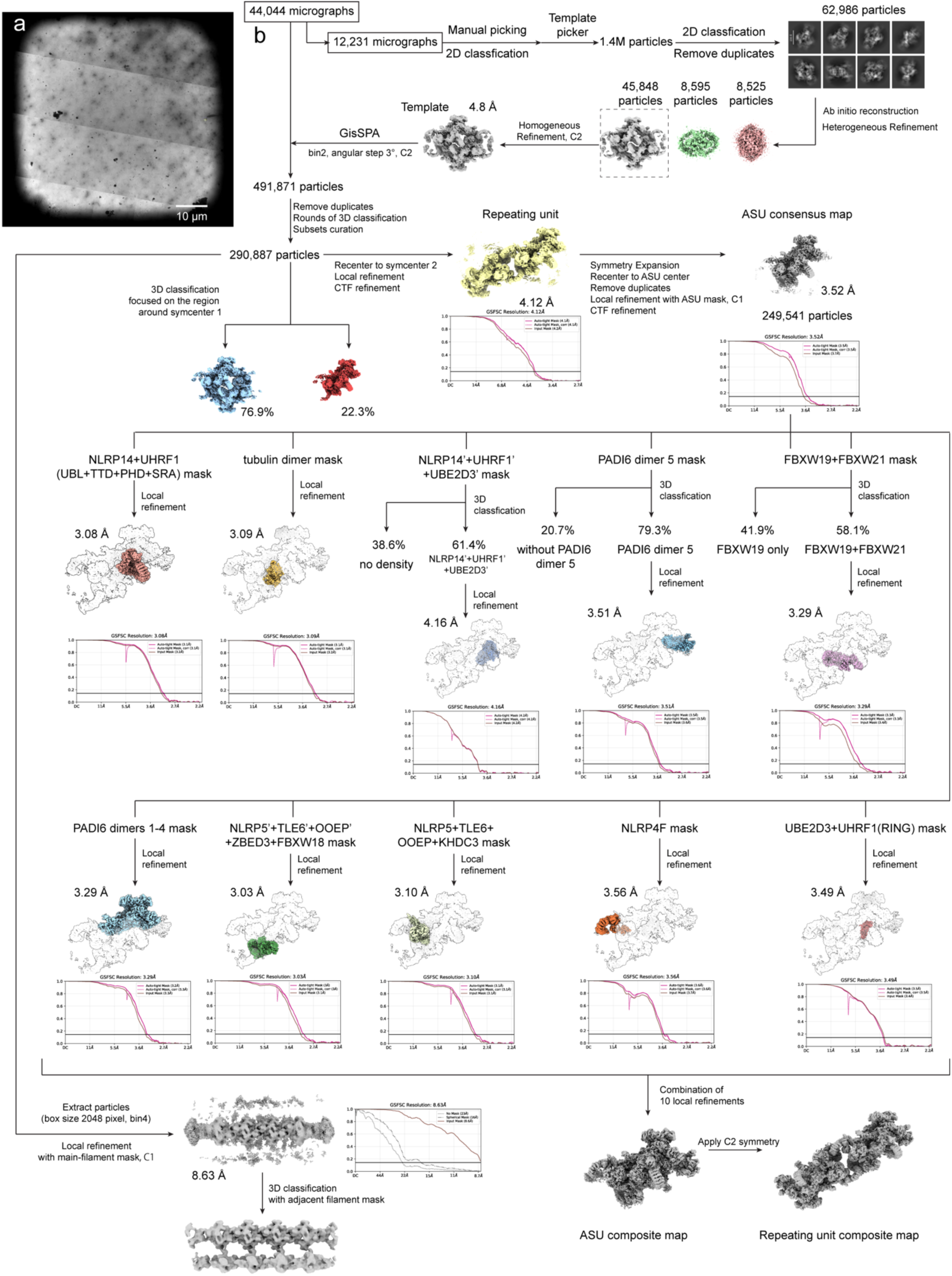
Cryo-EM sample preparation and single-particle data processing workflow for the native CPL. **a**, Representative intermediate-magnification cryo-EM micrograph. **b**, Single-particle cryo-EM data processing workflow.

**Extended Data Fig. 3.**
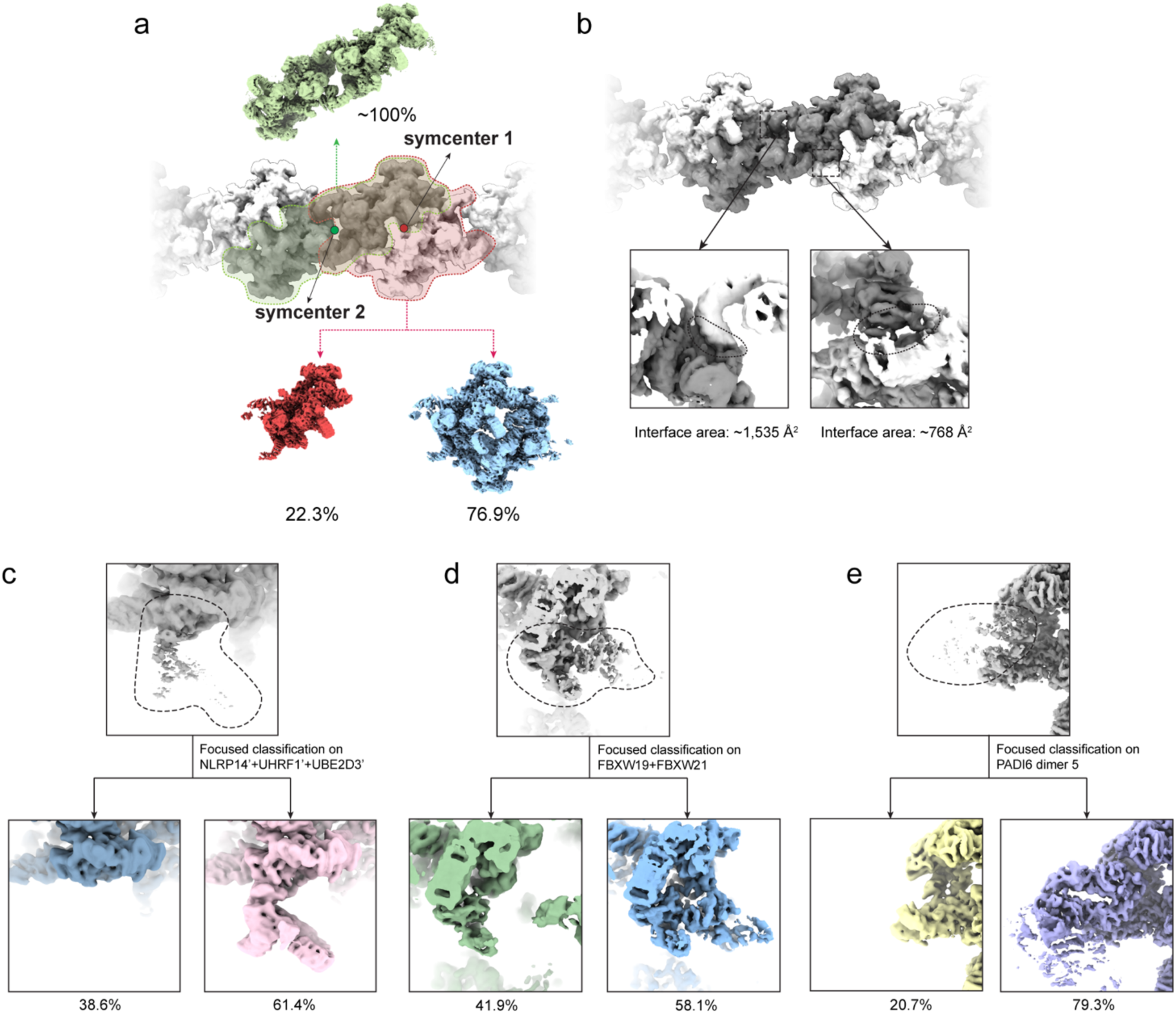
Focused classification analyses during CPL structure determination. **a,** Classification results for defining the biologically relevant CPL repeating unit. Two local twofold symmetry axes (symcenter 1 and symcenter 2) can be defined along the filament, and focused classification was performed on both candidate C2 assembly units. **b,** Contact-surface analysis of the two alternative repeating-unit definitions. The symcenter-2 assembly exhibits a doubled buried interface size. **c-e**, Focused classifications performed on different regions of the CPL map. Percentages indicate the fraction of particles assigned to each class.

**Extended Data Figure 4.**
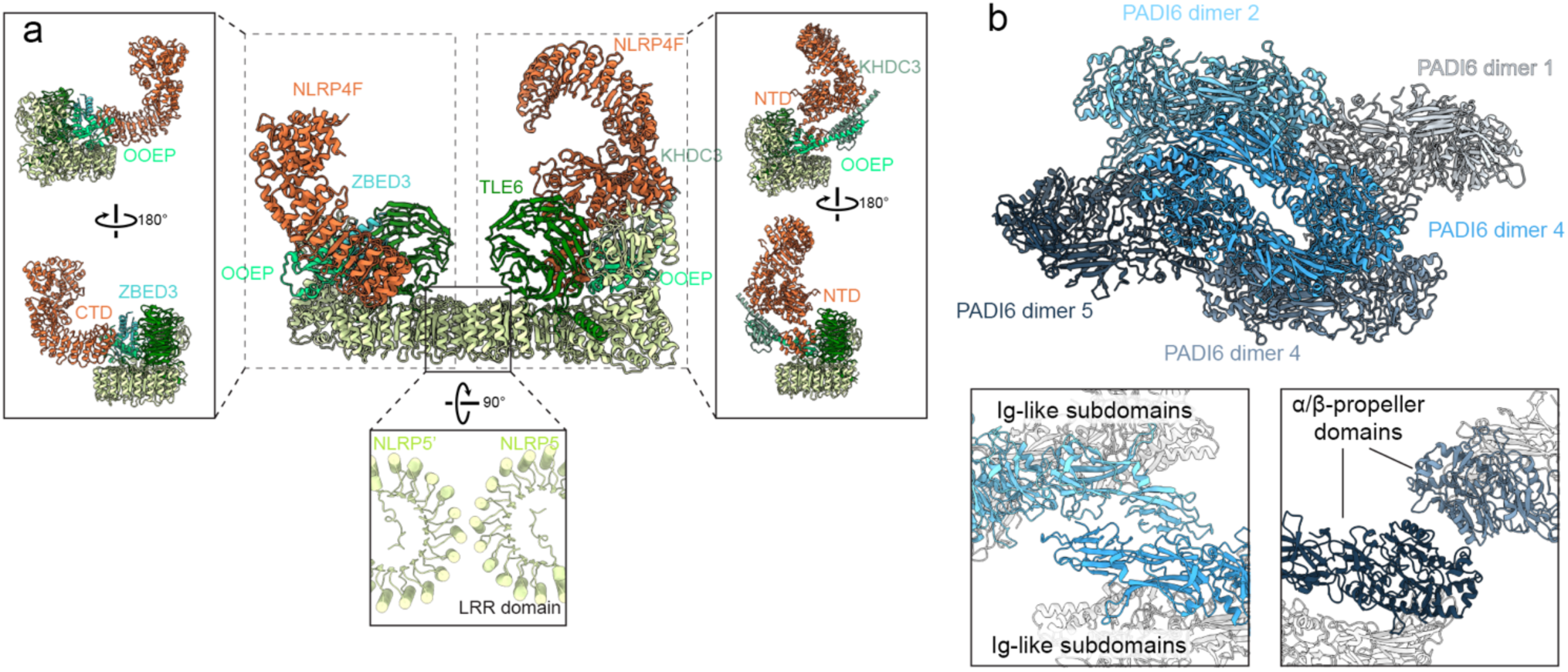
Structural organization of the CPL maternal-effect protein scaffold. **a**, The maternal-effect protein core forms an asymmetric heterodimer composed of NLRP5, TLE6, OOEP, ZBED3, KHDC3, and NLRP4F. Insets show side views of the two halves and the interface between the two NLRP5 LRR domains. The NLRP4F N-terminal domain (NTD) and C-terminal domain (CTD) are engaged in distinct binding modes in the two halves. **b**, Five PADI6 homodimers form a pentameric scaffold within each ASU. Insets highlight the inter-dimer contacts mediated by the N-terminal Ig-like subdomains and the C-terminal α/β-propeller domains.

**Extended Data Fig. 5.**
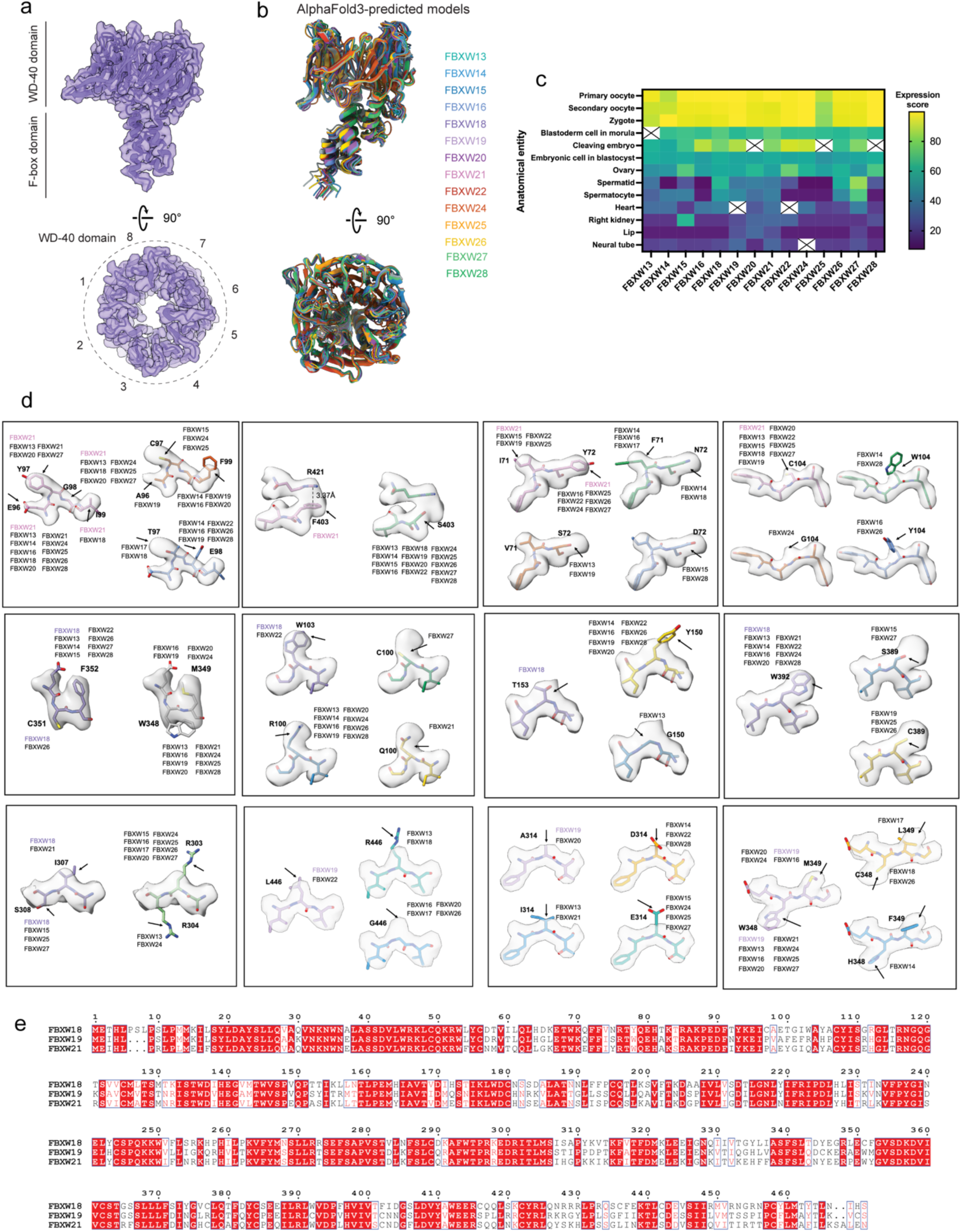
Assignment of CPL-associated FBXW proteins. **a,** Overall architecture of a representative CPL-associated FBXW protein, showing the N-terminal F-box domain and the C-terminal WD40 domain. Bottom, top view of the eight-bladed WD40 β-propeller. **b,** Overlay of AlphaFold3-predicted models for closely related FBXW-family members, illustrating their overall structural similarity. **c,** Expression heat map of candidate FBXW-family genes across developmental stages and tissues. Expression scores were retrieved from the Bgee database. **d,** Representative side-chain densities used to distinguish candidate FBXW-family members. Residues shown correspond to positions that differ among highly similar family members and enabled confident assignment of the three CPL-associated FBXW proteins. **e,** Sequence alignment of FBXW18, FBXW19, and FBXW21.

**Extended Data Fig. 6.**
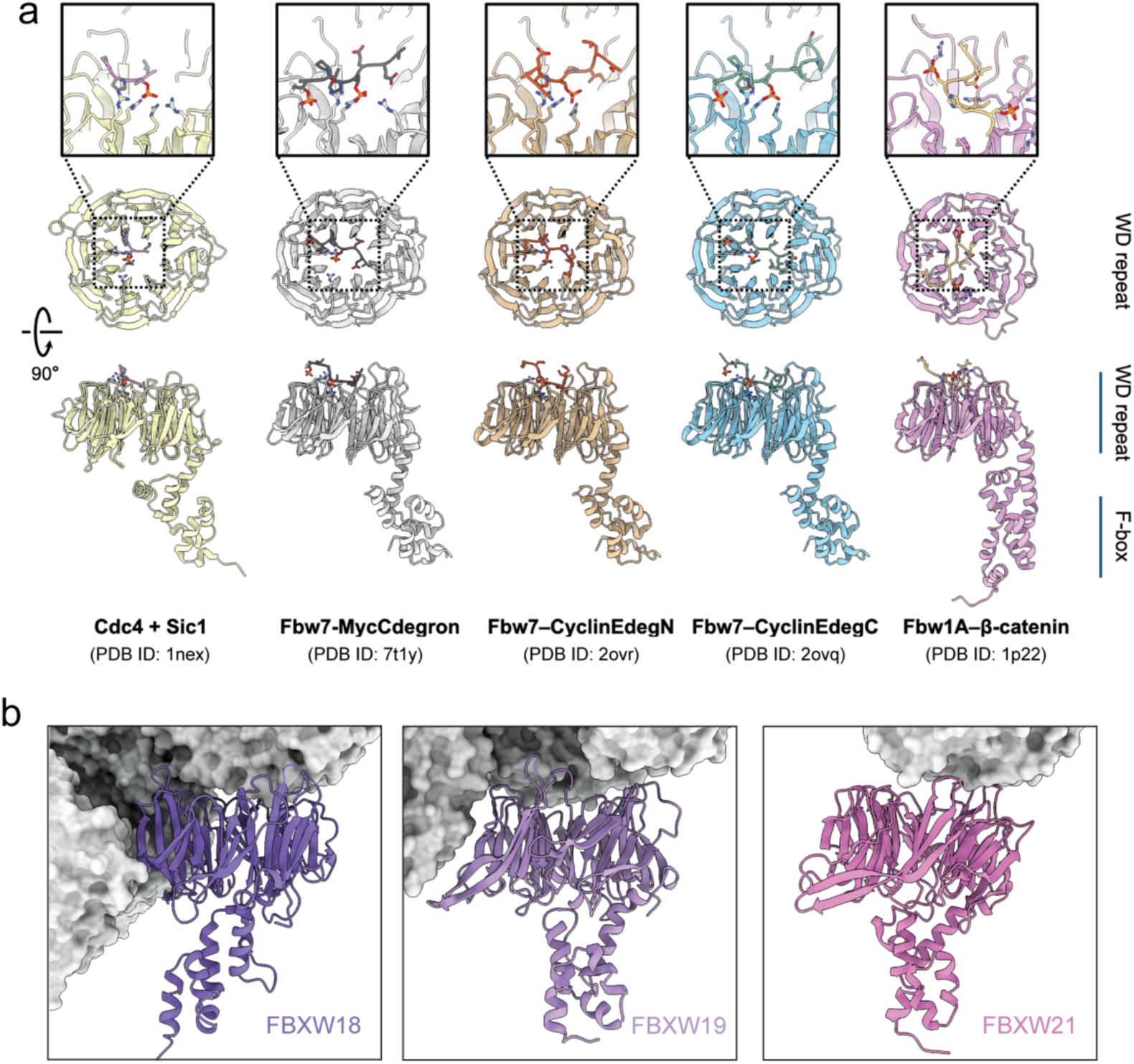
Canonical substrate-binding surfaces of FBXW proteins are buried in the CPL. **a,** Representative structures of substrate-bound FBXW-family proteins. In these complexes, substrates bind to the top surface of the WD40 β-propeller, centered around the pore. **b,** Structures of the three CPL-associated FBXW proteins (ribbons) positioned within the CPL assembly (gray surfaces). In all three cases, the top surfaces of the WD40 domain faces are largely buried.

**Extended Data Fig. 7.**
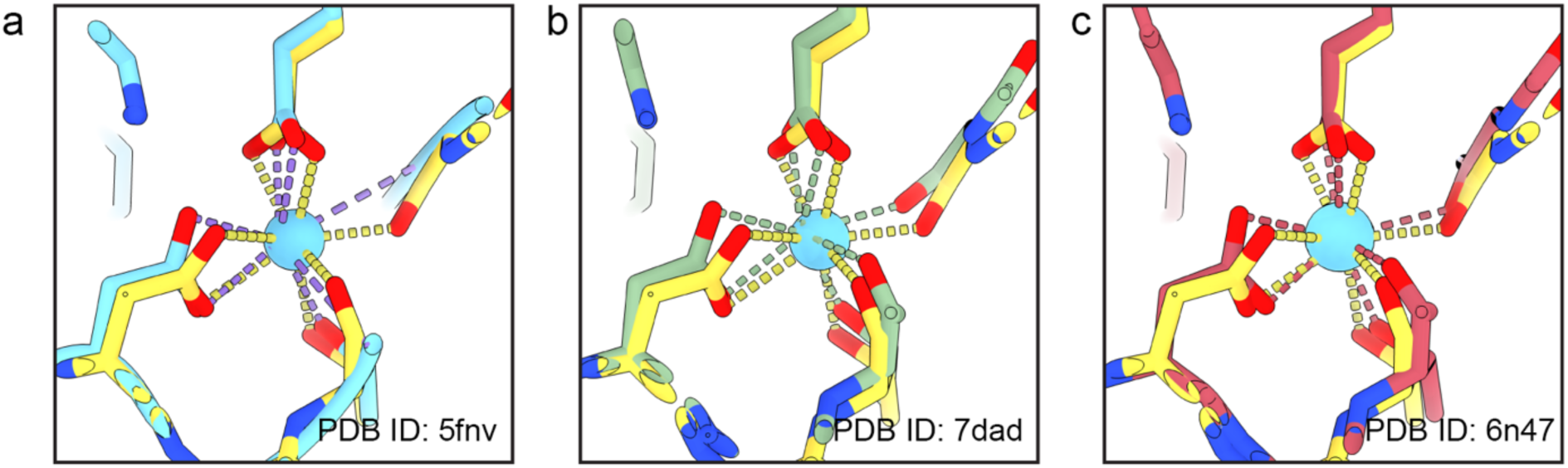
Comparison of the αK40-loop Ca²⁺ site with previously reported crystal structures of tubulin. a–c, Comparison of the current structure (yellow) with three reported structures (PDB IDs 5FNV (cyan), 7DAD (olive) and 6N47 (red)). In each case, the Ca²⁺ ion is shown as a cyan sphere and coordinating interactions are indicated by dashed lines.

**Extended Data Figure 8.**
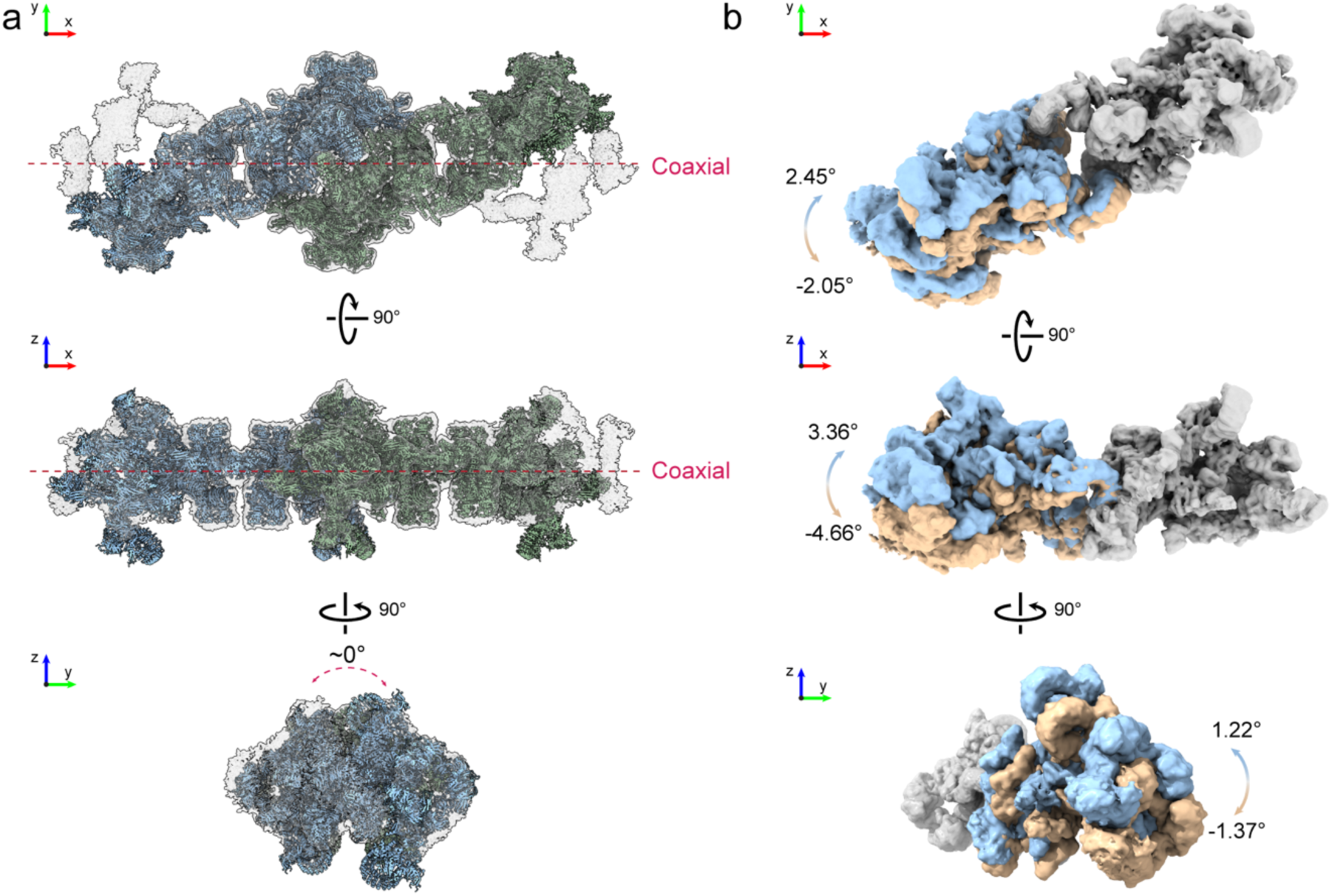
Structural analysis of intra-filament organization and flexibility. **a**, Model docking reveals that consecutive units align in a strictly coaxial manner with an average of ∼0° longitudinal rotation. **b**, The swinging of one ASU relative to the other across three dimensions reveals intra-unit conformational flexibility.

**Extended Data Table 1.** Data collection and refinement statistics.

|  | CPL ASU<br>(EMD-76334)<br>(PDB<br>ID:12DL) | CPL ASU<br>consensus<br>(EMD-76335) | CPL filament<br>(EMD-76333) | Local refine<br>(EMD-76315) | Local refine<br>(EMD-76321) | Local refine<br>(EMD-76322) |
| --- | --- | --- | --- | --- | --- | --- |
| **Data collection and processing** |  |  |  |  |  |  |
| Magnification | 81,000 | 81,000 | 81,000 | 81,000 | 81,000 | 81,000 |
| Voltage (kV) | 300 | 300 | 300 | 300 | 300 | 300 |
| Electron exposure (e-/ Å<sup>2</sup>) | 50 | 50 | 50 | 50 | 50 | 50 |
| Defocus range (µm) | -1.2 to -2.5 | -1.2 to -2.5 | -1.2 to -2.5 | -1.2 to -2.5 | -1.2 to -2.5 | -1.2 to -2.5 |
| Pixel size (Å) | 1.068 | 1.068 | 1.068 | 1.068 | 1.068 | 1.068 |
| Final Micrographs (no.) | 44,044 | 44,044 | 44,044 | 44,044 | 44,044 | 44,044 |
| Initial particle images (no.) | 491,871 | 491,871 | 491,871 | 491,871 | 491,871 | 491,871 |
| Final particle images (no.) | 249,541 | 249,541 | 290,887 | 249,541 | 249,541 | 249,541 |
| Symmetry imposed | C1 | C1 | C1 | C1 | C1 | C1 |
| Map resolution (Å) | 3.52 | 3.52 | 8.63 | 3.29 | 3.10 | 3.03 |
| FSC threshold | 0.143 | 0.143 | 0.143 | 0.143 | 0.143 | 0.143 |
| **Model Refinement** |  |  |  |  |  |  |
| Model resolution (Å) | 3.3 |  |  |  |  |  |
| FSC threshold | 0.5 |  |  |  |  |  |
| Model vs. map correlation<br>coefficient (cc_mask) | 0.83 |  |  |  |  |  |
| Model composition |  |  |  |  |  |  |
| Non-hydrogen atoms | 133305 |  |  |  |  |  |
| Protein residues | 16740 |  |  |  |  |  |
| Nucleotide | 0 |  |  |  |  |  |
| Ligands | 16 |  |  |  |  |  |
| B factors (Å<sup>2</sup>) |  |  |  |  |  |  |
| Protein | 118 |  |  |  |  |  |
| Nucleotide |  |  |  |  |  |  |
| Ligand | 84 |  |  |  |  |  |
| R.m.s. deviations |  |  |  |  |  |  |
| Bond lengths (Å) | 0.04 |  |  |  |  |  |
| Bond angles (°) | 0.6 |  |  |  |  |  |
| Validation |  |  |  |  |  |  |
| MolProbity score | 1.5 |  |  |  |  |  |
| Clashscore | 4.1 |  |  |  |  |  |
| Poor rotamers (%) | 0 |  |  |  |  |  |
| Ramachandran plot |  |  |  |  |  |  |
| Favored (%) | 95.03 |  |  |  |  |  |
| Allowed (%) | 4.88 |  |  |  |  |  |
| Disallowed (%) | 0.09 |  |  |  |  |  |

**Extended Data Table 2.** Data collection and refinement statistics.

|  | Local refine<br>(EMD-763223) | Local refine<br>(EMD-76324) | Local refine<br>(EMD-76325) | Local refine<br>(EMD-76326) | Local refine<br>(EMD-76327) | Local refine<br>(EMD-76330) | Local refine<br>(EMD-76331) |
| --- | --- | --- | --- | --- | --- | --- | --- |
| **Data collection and processing** |  |  |  |  |  |  |  |
| Magnification | 81,000 | 81,000 | 81,000 | 81,000 | 81,000 | 81,000 | 81,000 |
| Voltage (kV) | 300 | 300 | 300 | 300 | 300 |  |  |
| Electron exposure (e-/ Å²) | 50 | 50 | 50 | 50 | 50 | 50 | 50 |
| Defocus range (µm) | -1.2 to -2.5 | -1.2 to -2.5 | -1.2 to -2.5 | -1.2 to -2.5 | -1.2 to -2.5 | -1.2 to -2.5 | -1.2 to -2.5 |
| Pixel size (Å) | 1.068 | 1.068 | 1.068 | 1.068 | 1.068 | 1.068 | 1.068 |
| Final Micrographs (no.) | 44,044 | 44,044 | 44,044 | 44,044 | 44,044 | 44,044 | 44,044 |
| Initial particle images (no.) | 491,871 | 491,871 | 491,871 | 491,871 | 491,871 | 491,871 | 491,871 |
| Final particle images (no.) | 249,541 | 249,541 | 249,541 | 144,983 | 249,541 | 197,886 | 154,218 |
| Symmetry imposed | C1 | C1 | C1 | C1 | C1 | C1 | C1 |
| Map resolution (Å) | 3.08 | 3.08 | 3.56 | 3.29 | 3.49 | 3.51 | 4.16 |
| FSC threshold | 0.143 | 0.143 | 0.143 | 0.143 | 0.143 | 0.143 | 0.143 |

