## Supplementary Note 1 for "The cytoplasmic lattice in mammalian eggs sequesters ubiquitination machinery and tubulin in reserve"

**Supplementary Note 1 | Estimation of the fraction of tubulin associated with the CPL**

The quantitative values used in these calculations were obtained from tandem mass tag (TMT)-based quantitative proteomics, where protein abundance is reported as summed reporter ion intensities (area) for each protein across all assigned peptides. These area values reflect the integrated signal derived from TMT reporter ions and are proportional to the relative abundance of each protein in the sample.

From this dataset, the measured abundance (area) values for CPL components were:

NLRP5 = 6.12 × 10⁹
TLE6 = 6.30 × 10⁹
OOEP = 7.18 × 10⁹

CPL-associated tubulin was estimated using the stoichiometry derived from the structure, in which NLRP5, TLE6, and OOEP are present at 1:1 ratios with tubulin heterodimers, with two copies of each protein per structural unit:

CPL tubulin (NLRP5) = (6.12 × 10⁹) / 2 × 2 = 6.12 × 10⁹
CPL tubulin (TLE6) = (6.30 × 10⁹) / 2 × 2 = 6.30 × 10⁹
CPL tubulin (OOEP) = (7.18 × 10⁹) / 2 × 2 = 7.18 × 10⁹

Total tubulin abundance was calculated independently by summing the area values of all detected α- and β-tubulin isoforms in the dataset:

Total tubulin (monomers) = 4.90 × 10¹⁰

The fraction of tubulin associated with the CPL was then calculated as:

CPL-bound fraction (NLRP5) = (6.12 × 10⁹) / (4.90 × 10¹⁰) = 0.125 ≈ 12.5%
CPL-bound fraction (TLE6) = (6.30 × 10⁹) / (4.90 × 10¹⁰) = 0.129 ≈ 12.9%
CPL-bound fraction (OOEP) = (7.18 × 10⁹) / (4.90 × 10¹⁰) = 0.147 ≈ 14.7%
